# Regulated resource reallocation is transcriptionally hard wired into the yeast stress response

**DOI:** 10.1101/2024.12.03.626567

**Authors:** Rachel A. Kocik, Audrey P. Gasch

## Abstract

Many organisms maintain generalized stress responses activated by adverse conditions. Although details vary, a common theme is the redirection of transcriptional and translational capacity away from growth-promoting genes and toward defense genes. Yet the precise roles of these coupled programs are difficult to dissect. Here we investigated *Saccharomyces cerevisiae* responding to salt as a model stressor. We used molecular, genomic, and single-cell microfluidic methods to examine the interplay between transcription factors Msn2 and Msn4 that induce stress-defense genes and Dot6 and Tod6 that transiently repress growth-promoting genes during stress. Surprisingly, loss of Dot6/Tod6 led to slower acclimation to salt, whereas loss of Msn2/4 produced faster growth during stress. This supports a model where transient repression of growth-promoting genes accelerates the Msn2/4 response, which is essential for acquisition of subsequent peroxide tolerance. Remarkably, we find that Msn2/4 regulate *DOT6* mRNA production, influence Dot6 activation dynamics, and are required for full repression of growth-promoting genes. Thus, Msn2/4 directly regulate resource reallocation needed to mount their own response. We discuss broader implications for common stress responses across organisms.

**SYNOPSIS:** This study investigates how genes induced and repressed in the yeast Environmental Stress Response contribute to stress tolerance, growth rate, and resource allocation. The work uses molecular, genomic, and systems biology approaches to present new insights into eukaryotic responses to acute stress.

**HIGHLIGHTS:**

- Cells lacking stress-activated transcription factors have a faster post-stress growth rate
- Cells lacking repressors of growth-promoting genes have a slower post-stress growth rate
- Stress-defense factors control the induction of growth-promoting gene repressors, thereby coordinating the resource re-allocation needed for the response

## INTRODUCTION

Cells have evolved intricate systems to allocate limited intracellular resources according to the demands of an often-fluctuating environment. When conditions are favorable and nutrients are plentiful, many microbes maximize their growth rate by directing resources toward processes required for growth and proliferation. Much of the transcriptional and translational capacity goes toward producing ribosomes, which under optimal conditions fuel rapid growth (Scott & Hwa, 2011, 2023; Warner, 1999). However, in suboptimal conditions, especially in response to acute stress, resources including transcriptional and translational capacity are reallocated toward survival, often at the expense of growth and growth-promoting processes. In fact, rapid growth and high stress tolerance represent a well-known tradeoff: Fast growing cells directing resources toward division are often the most sensitive to stress, whereas slow growing cells are typically highly stress tolerant (Balaban et al., 2004; Basu et al., 2022; Levy et al., 2012; Pontes & Groisman, 2019; Zakrzewska et al., 2011; Zhang et al., 2020). This is true across organisms, including bacteria, yeast, plants, and mammalian cells. However, it remains poorly understood how cells regulate changes in resource allocation during times of stress and which cellular objectives (maximal growth versus high stress tolerance) dictate those changes. This is important for understanding how cells thrive in natural environments that are dynamic and often sub-optimal. Presumably, cells must coordinate multiple facets of biology as they respond and acclimate to changing conditions.

Budding yeast *Saccharomyces cerevisiae* has been an excellent model to understand fundamental principles of growth-versus-defense responses. Upon an acute shift to suboptimal conditions, yeast activate condition-specific responses customized for each condition, along with a common transcriptomic response known as the environmental stress response (ESR) (Causton et al., 2001; Gasch et al., 2000). The ESR is activated in response to diverse types of stress, including nutrient limitation, shifts in environmental conditions like osmolarity or temperature, and exposure to toxic compounds. The program includes ∼300 transcriptionally induced genes (iESR genes) involved in wide-ranging defense processes such as redox balance, protein folding and degradation, carbohydrate and energy metabolism, trehalose and glycogen biosynthesis, and other processes (Gasch, 2002a; Gasch et al., 2000). Induced transcription of the iESR genes is coordinated with reduced expression of ∼600 genes (rESR genes) that encode ribosomal proteins (RP) and proteins involved ribosome biogenesis, translation, and other growth-promoting processes (RiBi genes) (Gasch, 2002a; Gasch et al., 2000). In optimal conditions, cells devote significant resources to transcribing and translating rESR transcripts, which are required to promote rapid growth, while maintaining low production of iESR and defense proteins (**Fig 1A**). Notably, other organisms maintain analogous, if not orthologous, responses to balance stress-defense versus growth-promoting processes, such as the Integrated Stress Response (ISR) in mammals (Costa-Mattioli & Walter, 2020; Harding et al., 2003) and the Stringent / SOS responses in bacteria (Gourse et al., 2018; Irving et al., 2021). Despite differences in the regulation of these programs across species, many of the underlying themes are conserved, including the redirection of translational capacity toward stress-induced transcripts. Yet decoupling the role of translational suppression from the functions of induced proteins has remained challenging.

**Figure 1.**
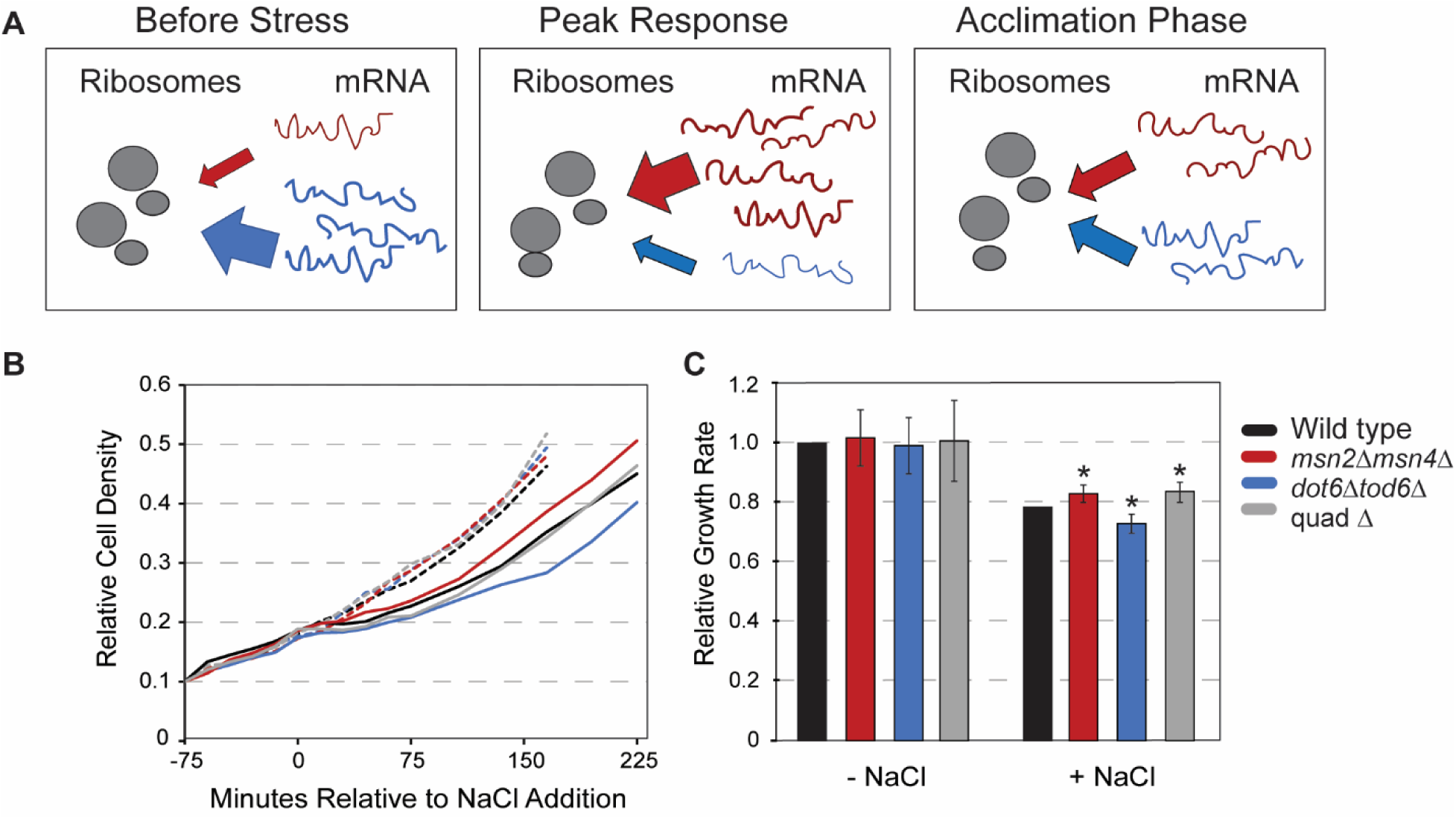
Msn2/4 versus Dot6/Tod6 have opposing influences on post-stress growth rate. **A)** A model for the dynamics of stress-defense (red) mRNAs induced by stress versus growth-promoting (blue) mRNAs that are highly expressed in the absence of stress but transiently repressed during stress acclimation. **B)** Representative relative cell density (OD_600_) for wild-type (black), *msn2Δmsn4Δ* (red), *dot6Δtod6Δ* (blue), and quadruple mutant (“quadΔ”) (grey) growing in the absence (dashed lines) and presence (solid lines) of 0.7 M NaCl added at 0 minutes. **C)** Average and standard deviation (n=7) of growth rates in the absence (left) and presence (right) of 0.7 M NaCl (calculated from 75 to 225 min timepoints). Post NaCl growth rates were calculated relative to their paired wild-type, then scaled to the average wild-type post-stress versus pre-stress relative rate. *, p-value < 0.01, two-tailed, replicate-paired t-test relative to the correspondingly treated wild-type.

iESR induction and rESR repression are highly correlated in bulk cultures, at least in part due to coordinated regulation (Gasch, 2002a). iESR gene induction is coordinated by the paralogous ‘general stress’ transcription factors Msn2 and Msn4 (Causton et al., 2001; Estruch & Carlson, 1993; Gasch et al., 2000), overlaid with condition-specific factors that customize expression levels in specific environments (Gasch, 2002a). Msn2 and/or Msn4 (Msn2/4) bind the stress response promoter element (STRE, CCCCT) present in one to many copies upstream of hundreds of target genes (Martínez-Pastor et al., 1996; Stewart-Ornstein et al., 2013). Like many stress-regulated factors, Msn2/4 are regulated by nuclear translocation: phosphorylation at specific residues, including by growth-promoting Protein Kinase A (PKA) and TOR kinases, restricts the factors to the cytoplasm during optimal conditions (Beck & Hall, 1999; Boy-Marcotte et al., 1998; Görner et al., 1998; Jacquet et al., 2003; Smith et al., 1998). During stress, dephosphorylation of those residues coupled with other mechanisms prompts Msn2/4 nuclear localization and gene induction (De Wever et al., 2005; Garreau et al., 2000; González et al., 2009; Görner et al., 1998; Lenssen et al., 2005; Santhanam et al., 2004; Smith et al., 1998). The dynamics of nuclear translocation can impart important differences in gene expression, depending on the amount of protein that enters the nucleus, the duration of nuclear accumulation, and gene-promoter architecture including the number of upstream STRE elements of different genes (Gasch, 2002a; Hansen & O’Shea, 2013, 2015b, 2015a, 2016; Hansen & Zechner, 2021; Hao & O’Shea, 2012; Purvis et al., 2012; Purvis & Lahav, 2013; Stewart-Ornstein et al., 2013; Sweeney & McClean, 2023). rESR subgroups are modulated by different regulators (Gasch, 2002a; Jorgensen et al., 2004; Marion et al., 2004; Schawalder et al., 2004; Shore & Nasmyth, 1987). Many of the RiBi genes are repressed during stress by Dot6 and its paralog Tod6 (Bergenholm et al., 2018; Cheng & Brar, 2019; Lippman & Broach, 2009). Dot6 and/or Tod6 (Dot6/Tod6) bind the GATGAG motif, which is present in about ∼70% of RiBi gene promoters (Badis et al., 2008; C. Zhu et al., 2009). Furthermore, Dot6/Tod6 are also regulated by nuclear translocation, where phosphorylation by PKA and Sch9/TOR maintains the factors in cytoplasm and dephosphorylation leads to nuclear localization (Huber et al., 2011; Lippman & Broach, 2009).

During many stress responses, ESR activation, and in particular rESR repression, coincides with growth reduction; however, the ESR is not an indirect response to growth as previously proposed (see (Brauer et al., 2008; Castrillo et al., 2007; Lee et al., 2011; Lu et al., 2009; O’Duibhir et al., 2014; Regenberg et al., 2006)). Cells already arrested in division and with reduced biomass production still show rESR repression upon stress exposure (Ho et al., 2018). We proposed that rESR repression during acute stress helps to redirect transcriptional and translational capacity toward induced mRNAs (Bergen et al., 2022; Ho et al., 2018; Lee et al., 2011). Cells lacking *DOT6/TOD6* fail to fully repress rESR genes during acute salt stress, leading to the over-abundance of rESR transcripts, which remain associated with ribosomes. In turn, iESR transcripts show reduced ribosome binding and a delay in the production of iESR proteins (Ho et al., 2018; Lee et al., 2011). Indeed, the *dot6Δtod6Δ* mutant also shows delayed synthesis of iESR protein Ctt1 encoding cytosolic catalase, despite higher induction of *CTT1* mRNA (Bergen et al., 2022; Ho et al., 2018). Thus, repression of the rESR genes likely serves to indirectly accelerate the stress response by removing rESR mRNAs from the translating pool.

Studying cell cultures in bulk can obscure causal relationships that vary across individual cells in a population. Thus, investigating cell-to-cell heterogeneity has been a useful tool in deciphering co-varying phenotypes that can reflect on cellular coordination (Bagamery et al., 2020; Barber et al., 2021; Gasch et al., 2017; Levy et al., 2012; Li et al., 2018). We previously used microfluidic live-cell imaging to study single-cell heterogeneity in cells responding to an acute dose of sodium chloride (NaCl) stress (Bergen et al., 2022). Our system enabled characterization of multiple phenotypes in single cells, including growth rate, colony size, cell-cycle phase and nuclear translocation dynamics of fluorescently tagged Msn2-GFP and Dot6-mCherry expressed in the same cells. Somewhat counterintuitively, we found that wild-type cells with larger Dot6 nuclear translocation response, predicted to produce stronger repression of growth-promoting rESR genes, in fact acclimate with faster post-stress growth rates following acute salt stress. Wild-type cells with stronger Dot6 activation also displayed faster production of Ctt1 protein compared to cells with weaker activation, consistent with our model that rESR repression helps to accelerate production of induced proteins (Bergen et al., 2022). We proposed that Dot6-dependent transcriptional repression, and by extension repression of the rESR as a whole, is important to reallocate resources for faster acclimation to stress conditions. But if and how a faster response is important for stress survival has not been tested. Furthermore, how resource reallocation through gene repression is coordinated with genes induced in the ESR was not clear from past work.

Here we investigated the interplay between Msn2/4 and Dot6/Tod6 activation dynamics, transcriptional regulation, and growth versus defense objectives. We confirmed that Dot6/Tod6 are required for normal acclimation dynamics following salt stress. Somewhat surprisingly, Msn2/4 are not only dispensable for salt survival but in fact come with a significant cost: cells lacking *MSN2/4* grow substantially faster than wild-type cells during salt acclimation. Consistent with prior work from our lab, Msn2/4 are essential for salt-induced acquisition of subsequent peroxide tolerance in a phenomenon known as acquired stress resistance (Berry et al., 2011; Berry & Gasch, 2008). Dot6/Tod6 also contribute to acquired peroxide tolerance by accelerating acquisition of peroxide tolerance. Strikingly, we discovered through microfluidics and RNA-seq transcriptomics that Msn2/4 regulate the abundance and activation of Dot6, and directly contribute to the salt-dependent repression of rESR genes. Thus, Msn2/4 help to direct resources toward its own program by modulating Dot6 behavior and rESR repression. We discuss implications of these results in the context of bacterial and mammalian stress responses for a unified framework for coordinated resource management during adversity.

## RESULTS

### Msn2/4 versus Dot6/Tod6 have opposing influences on post-stress growth rate

We previously showed that cells lacking Dot6 and Tod6 (*dot6Δtod6Δ)* grow indistinguishably to wild-type in the absence of stress but display slower growth after salt stress (Bergen et al., 2022). We confirmed here that *dot6Δtod6Δ* cells show a similar lag phase but reduced growth rate compared to wild-type cells only after salt treatment (p=0.002, replicate-paired T-test) (**Fig 1B**). A remaining question was the role of Msn2/4 and iESR induction on stress acclimation. We observed that, like the *dot6Δtod6Δ* mutant, *msn2Δmsn4Δ* cells grew indistinguishably to wild-type in the absence of stress (**Fig 1B**). Surprisingly, however, the *msn2Δmsn4Δ* mutant grew significantly faster than wild-type cells after salt stress (p = 0.008). These cells also appeared to show a shorter lag, since they had a significantly greater percent change in cell density over 60 minutes compared to the wild-type (p = 0.010, replicate and time-paired two-tailed T-test). We conclude that mounting the iESR comes with a significant cost to post-stress growth rate; this also reveals that the main objective of cells during this phase is something other than maximizing growth, since cells are clearly capable of growing faster (see more below).

Given that *MSN2/4* deletion accelerates salt acclimation and the *DOT6/TOD6* deletion retards it, we wondered how deleting all four factors would influence growth. If Dot6/Tod6-dependent repression serves to release resources for induced protein production, then loss of the costly Msn2/4 response may recover growth rate in *dot6Δtod6Δ* cells. Alternatively, if Dot6 /Tod6 play a different unrecognized role, deletion of *MSN2/4* would not alleviate its post-stress growth requirement. We generated a strain lacking all four transcription factors (referred to as the quad mutant or *quadΔ*). Like both double mutants, the quad mutant grew similarly to wild-type in the absence of stress. Initially after salt treatment (over 30 – 60 minutes), the *quadΔ* change in cell density was similar to the wild-type and *dot6Δtod6Δ* strains (**Fig 1B**). However, as the cells continued to grow (75 – 225 minutes post-salt addition), the quad mutant grew faster than the wild-type, similar to cells lacking Msn2/4. Thus, the reduced post-stress growth rate of *dot6Δtod6Δ* cells acclimating to salt stress can be complemented by loss of Msn2/4 (see Discussion).

### Both Msn2/4 and Dot6/Tod6 responses provide a benefit to future-stress survival

The cost of Msn2/4 activity to post-stress growth rate raised questions about why cells would maintain this response. One explanation is acquired stress resistance, in which cells that mount a stress response during a mild dose of one stress can survive what would otherwise be a lethal dose of subsequent stress treatment (Berry & Gasch, 2008). Past work from our lab showed that Msn2/4 are essential for acquired resistance to severe peroxide stress after salt-stress pretreatment (Berry et al., 2011; Berry & Gasch, 2008), which we confirmed here. At varying times before and after NaCl treatment, an aliquot of culture was removed and cells were exposed to a panel of H_2_O_2_ doses for 2 hours, after which colony forming units were assessed (**Fig 2A**). The relative viability at each dose was normalized to the side-by-side treated wild type, and a single H_2_O_2_ survival score was calculated as the sum of those scores across doses (see Methods, **Fig EV1**). As expected, cells lacking *MSN2/4* had a major defect acquiring peroxide tolerance after salt treatment, as did the quad mutant (**Fig 2B**). The acquisition of peroxide tolerance was also dependent on Msn2/4 target *CTT1* encoding cytosolic catalase, since a strain lacking *CTT1* acquired no tolerance, as shown previously (Berry & Gasch, 2008, 2008; Guan et al., 2012) (**Fig 2B**). Thus, the Msn2/4 response, at least in part via Ctt1, is essential for acquired peroxide tolerance under these conditions.

**Figure 2.**
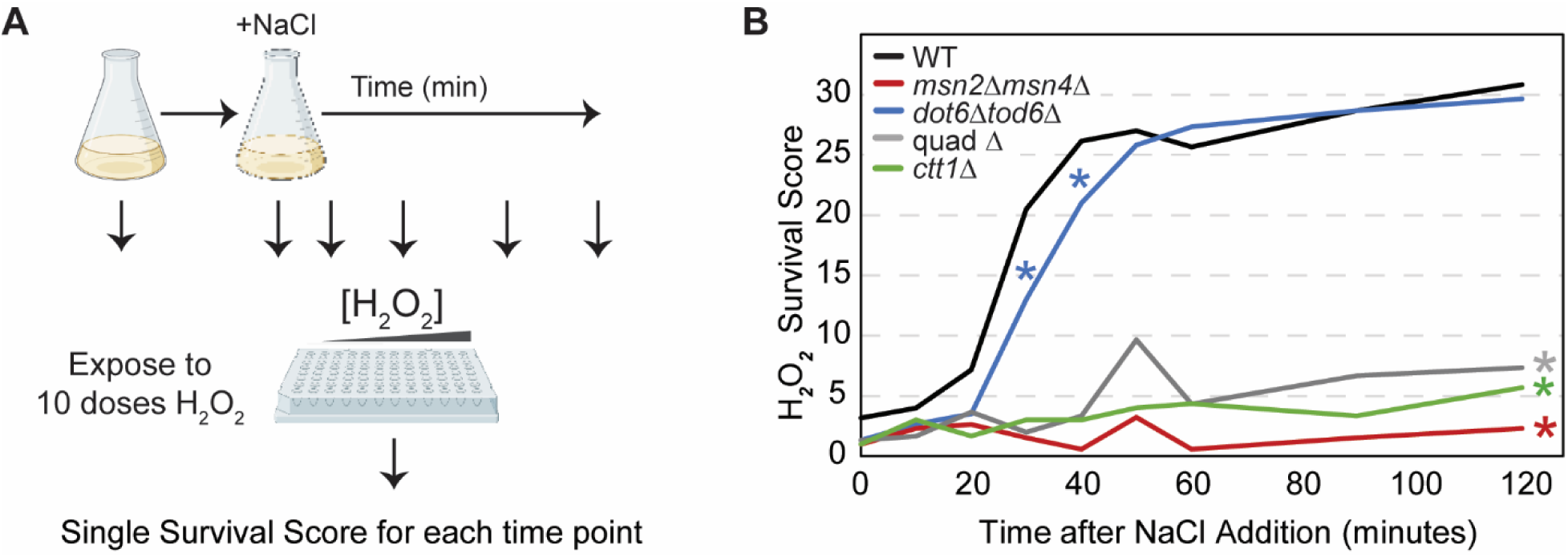
Msn2/4 and Dot6/Tod6 responses are important for acquired stress resistance. **A)** Schematic of acquired stress resistance protocol, see text. **B)** The average change in H_2_O_2_ survival scores for wild-type (black), *msn2Δmsn4Δ* (red), *dot6Δtod6Δ* (blue), quad Δ (grey), and *ctt1*Δ (green) cells. n=3 replicates except for *dot6Δtod6Δ* where n = 6. *, p-value < 0.03 (*), one-tailed, replicate-paired t-test at each timepoint (see **Fig EV1** for paired datasets used in statistics). The *msn2Δmsn4Δ, quadΔ and ctt1Δ* mutants were highly significant at all time points after 20 min (represented by a single asterisk at the end of the curve).

The role of Dot6/Tod6 and rESR repression in acquired stress resistance had not been previously investigated. Importantly, we found that Dot6/Tod6 were also required for normal acquisition of peroxide treatment. While the *dot6Δtod6Δ* mutant acquired wild-type levels of hydrogen peroxide resistance after exposure to salt, they did so with a significant delay (**Fig 2B**). Wild-type cells acquired maximal resistance by ∼40 min; however, the *dot6Δtod6Δ* cells took over ∼60 minutes to reach maximal tolerance. This delayed acquisition of peroxide tolerance cannot be explained by the reduced growth rate of the mutant, which was observed at later time points (see **Fig 1**). Instead, the timing of the delay correlates with delayed Ctt1 protein production in the *dot6Δtod6Δ* mutant (Bergen et al., 2022; Ho et al., 2018). These results are consistent with the model that Dot6/Tod6 gene repression helps to accelerate production of stress-defense proteins needed for acquired stress resistance.

### Interplay between Msn2 and Dot6 activation dynamics

While response couplings are often difficult to identify in bulk cultures, co-varying phenotypes become apparent when scoring single-cell heterogeneity within a population. We previously developed a microfluidics assay to explore heterogeneity in the nuclear translocation dynamics of Msn2-mCherry and Dot6-GFP expressed in the same cells (Bergen et al., 2022). In that study, we found that wild-type cells with a larger peak in Dot6 nuclear accumulation acclimated with faster post-stress growth rates than cells with a smaller peak. Here we investigated the interplay between Dot6 and Msn2 activation dynamics in the presence or absence of the opposing factors.

To explore this, we generated mutants in which one or the other set of paralogous transcription factors was deleted. One strain expressed Msn2-mCherry in the absence of *DOT6/TOD6* while another strain expressed Dot6-GFP in the absence of *MSN2/4*. We compared the response of Dot6-GFP or Msn2-mCherry in each mutant to wild-type cells that expressed both Msn2-mCherry and Dot6-GFP as well as a third constitutive fluorescence marker (Nhp6a-iRFP, see Methods). This enabled mixing each mutant with the wild-type and distinguishing strains based on Nhp6a-iRFP (see Methods), providing sensitive comparison of strain behaviors in the same microfluidics chamber.

Loss of *DOT6/TOD6* did not appreciably influence Msn2-mCherry nuclear dynamics (**Fig 3**). Mutant and wild-type cells displayed similar distributions of Msn2 localization dynamics and Msn2-mCherry localization peak heights (**Fig 3B-C**). Furthermore, clustering of the individual cells based on Msn2 dynamics revealed that the two cell types cluster together and are not distinguishable by gross differences in Msn2 behavior (**Fig 3A**). We conclude that the presence of Dot6 does not significantly impact the behavior of Msn2.

**Figure 3.**
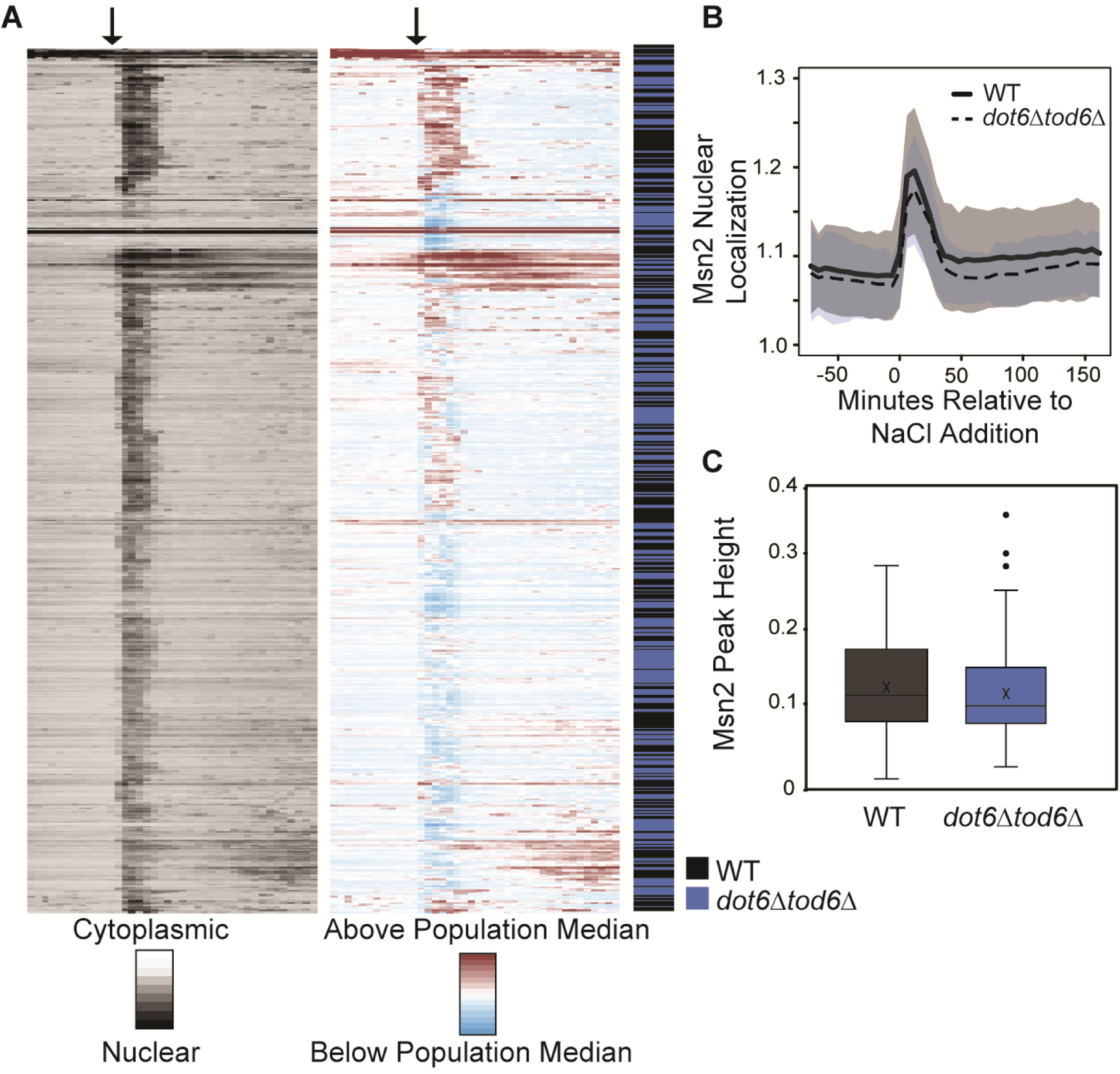
Msn2 behavior is not affected by a loss of Dot6 and Tod6. **A)** Left: Nuclear/cytoplasmic ratio was plotted for each cell (row) across timepoints (columns) before and after salt addition, indicated by the arrow. Right: the same data shown on the left normalized to the median of each column (population median). Cells were hierarchically clustered based on population-centered Msn2 nuclear translocation dynamics, after which cell identity was mapped onto the figure indicating wild-type (black) or *dot6Δtod6Δ* (blue) cells. **B)** The average Msn2 nuclear/cytoplasmic ratio in wild-type (black line) and *dot6Δtod6Δ* (dashed line) cells +/- one standard deviation. **C)** Distribution of Msn2 acute stress peak heights for wild-type (black) and *dot6Δtod6Δ* (blue) cells. p = 0.2, Wilcoxon rank-sum test.

In contrast, loss of Msn2/4 had a major impact on Dot6 behavior. Clustering cells based on Dot6-GFP translocation dynamics clearly delineated cell types: many of the *msn2Δmsn4Δ* cells had a weaker Dot6-GFP nuclear translocation response that was well below the median of all cells in the analysis, causing many of the *msn2Δmsn4Δ* cells to fall in a separate cluster (**Fig 4A**). The weaker response can also be seen in the distributions of nuclear translocation dynamics, where cells lacking *MSN2/4* displayed a lower Dot6 translocation response than the wild-type (**Fig 4B, EV2**). We noticed that, beyond just the translocation dynamics during acute stress, cells lacking *MSN2/4* showed significantly lower levels of Dot6-GFP signal overall, both before and after stress (**Fig 4C**). This cannot be explained by differences in cell size (which could change signal intensity over a changing area), since *msn2Δmsn4Δ* cell size is indistinguishable from wild-type both before or after salt stress (p = 0.7 and 0.2, respectively, Wilcoxon rank-sum test). Thus, cells lacking *MSN2/4* have less Dot6-GFP.

**Figure 4.**
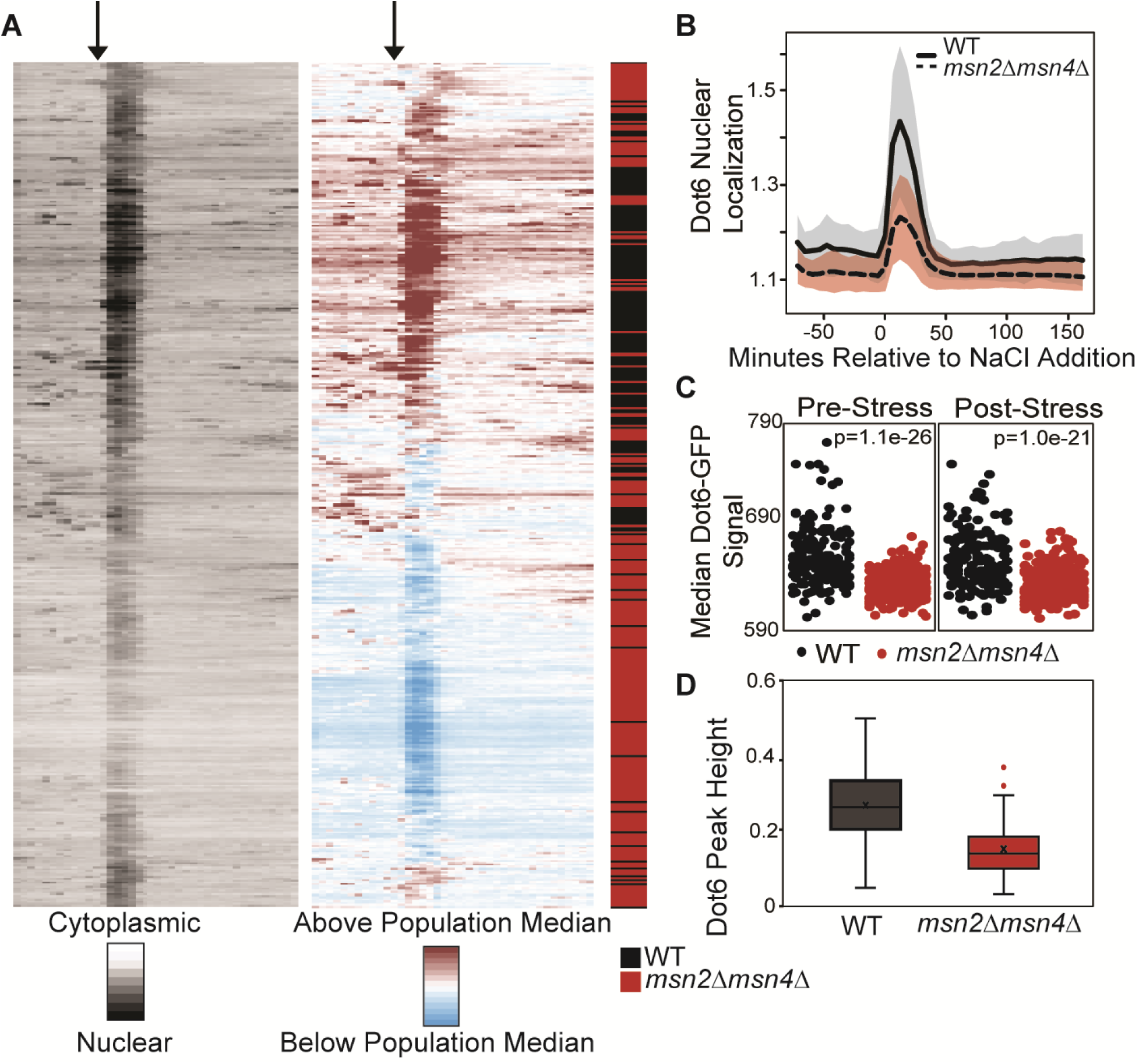
Loss of *MSN2/4* leads to decreased Dot6 abundance and nuclear localization. **A)** Wild type (WT) and *msn2Δmsn4Δ* cells from 4 replicates were clustered based on population-centered Dot6 nuclear translocation dynamics, as described in Fig 3. **B)** The population average of Dot6 nuclear/cytoplasmic ratio in wild-type (black line) and *msn2Δmsn4Δ* (dashed line) cells +/- one standard deviation. **C)** Distribution of median Dot6-GFP signal scored before (0-72 min) or after (120-216 min) NaCl treatment (see Methods for details) for WT and *msn2Δmsn4Δ* cells; p, Wilcoxon rank-sum test. **D)** Distribution of Dot6 acute-stress peak heights across 147 cells with similar Dot6-GFP levels, see Methods. p = 2.2e-10, Wilcoxon rank-sum test.

One question is if Msn2/4 has a direct impact on Dot6 levels/response or if the effect is indirect, perhaps simply due to the loss of iESR induction. We turned to past Msn2 chromatin-immunoprecipitation data from our lab to investigate (Huebert et al., 2012). Remarkably, both Msn2 and Msn4 bind the Dot6 promoter after multiple stress conditions in ours and other studies (Brodsky et al., 2020; Elfving et al., 2014; Huebert et al., 2012; Kuang et al., 2017; Ni et al., 2009) (**Fig EV3**). The Dot6 promoter harbors one perfect-match (CCCCT) to the Msn2 binding site at -580 bp and multiple other similar C-rich sequences within ∼600 bp upstream. These intriguing results suggest a direct conduit between Msn2 that contributes to iESR induction and regulation of Dot6 that participates in rESR repression.

We wondered if the apparent weaker nuclear translocation of Dot6-GFP in this mutant is artifactually influenced by having less GFP signal (see Methods). To test this, we investigated a subset of *msn2Δmsn4Δ* and wild-type cells who’s starting Dot6-GFP abundance was in the same range. Across 147 wild-type and *msn2Δmsn4Δ* cells with indistinguishable levels of Dot6-GFP (p = 0.5, Wilcoxon test, see Methods), the *msn2Δmsn4Δ* cells displayed significantly smaller Dot6-GFP nuclear translocation peak heights (**Fig 4D**, p = 2.2e-10, Wilcoxon test). Thus, cells lacking *MSN2/4* have lower Dot6 abundance overall, likely through direct regulation of *DOT6* transcription, and weaker Dot6 activation during salt stress, perhaps through indirect effects (see Discussion).

### Msn2/4 influence both iESR induction and rESR repression

A remaining question is if the weaker Dot6 activation in the *msn2Δmsn4Δ* mutant leads to weaker repression of Dot6 target genes. To directly address this question, we followed dynamic changes to the transcriptome before and in ten-minute increments after salt stress in the different strains. We first identified 489 genes whose response to salt treatment was altered in the *dot6Δtod6Δ* strain compared to wild-type (FDR < 0.05) in at least two timepoints (see Methods). 82% of these showed a repression defect, and of these 82% harbored upstream matches to the known Dot6 consensus (**Fig 5A**, p=3.3e-58 hypergeometric test). The remaining 18% of differentially expressed genes were induced by salt, but these generally showed subtle differences (and in some cases greater induction) compared to wild-type. These results indicate a primary role for Dot6/Tod6 in repression of their RiBi target genes.

**Figure 5.**
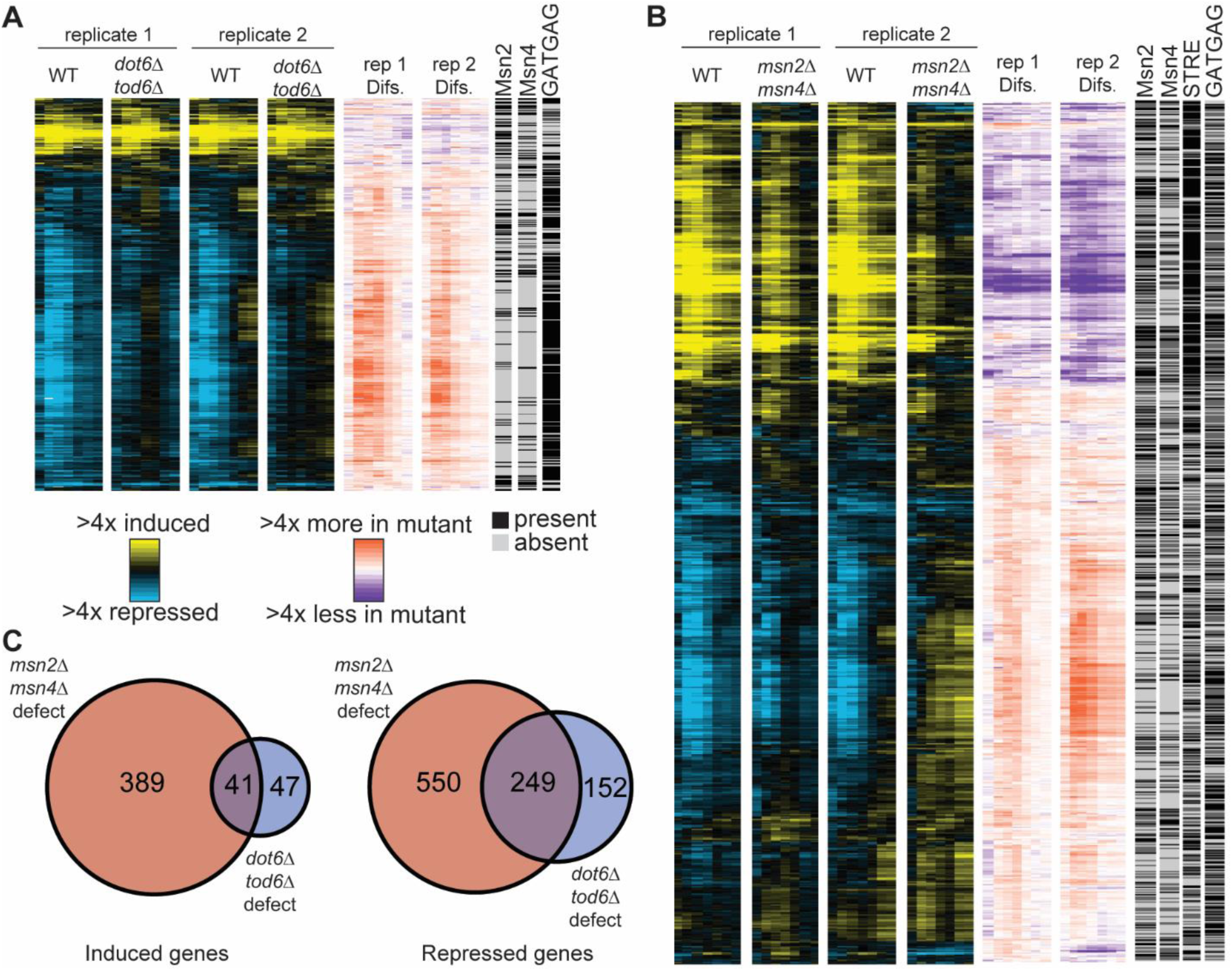
Msn2/4 influence iESR induction and rESR repression. **A)** 489 genes (rows) differentially expressed between wild-type and *dot6Δtod6Δ* cells (FDR < 0.05) in each timepoint (columns). Values represent log_2_(change) in expression compared to unstressed cells (blue-yellow plot) or the log_2_(difference) in fold-change values in *dot6Δtod6Δ* compared to wild-type cells, according to the keys. Columns on the right represent the presence (black) or absence (grey) of Msn2 binding, Msn4 binding, or Dot6 consensus (GATGAG) sequence (see Methods). **B)** As shown in A but for 1306 genes differentially expressed in *msn2Δmsn4Δ* compared to wild-type. **C)** Venn diagrams showing the overlap in induced or repressed genes with an induction or repression defect in the denoted mutant.

In contrast, loss of *MSN2/4* had wider ranging effects. We identified 1,306 differentially expressed genes in at least two timepoints in the *msn2Δmsn4Δ* strain compared to wild-type (FDR < 0.05). A third (430 genes) of these were induced by salt but with significantly weaker magnitude in *msn2Δmsn4Δ* cells (**Fig 5B**, orange/purple scale). This group was strongly enriched for genes whose promoters are bound by Msn2 and/or Msn4 and harbor upstream STRE elements (p=1.3e-24 and 6.7e-50, respectively, hypergeometric test), consistent with direct regulation. Of the remaining 63% of affected genes, many of these (799 genes) were repressed in wild-type cells but repressed to a lesser extent in the *msn2Δmsn4Δ* mutant. Two thirds (68%) of these genes contain upstream Dot6/Tod6 binding elements (p=3.7e-48, hypergeometric test) and 249 genes showed a repression defect at multiple time points in the *dot6Δtod6Δ* strain (**Fig 5C**). The weaker repression of these genes is consistent with weaker Dot6 activation in the absence of *MSN2/4*. However, we were surprised to find that many of these repressed genes also showed upstream Msn2/4 binding during stress: Of the genes whose salt-dependent repression required Msn2/4, 41% are bound by Msn2/4, which is more than expected by chance (p=3.0e-8, hypergeometric test) – however, unlike induced genes dependent on Msn2/4, the repressed genes were actually under-enriched for upstream STRE elements compared to chance (p=7e-8, hypergeometric test, **Fig 5B**). This result raises the possibility that Msn2/4 localization to rESR promoters occurs via a different mechanism than at iESR promoters.

We noted that 73% (181 genes) of the repressed genes that require both Msn2/4 and Dot6/Tod6 for full repression are devoid of upstream Msn2/4 binding during any stress condition. Together, these results suggest that Msn2/4 may play a more complex regulatory role than previously realized, affecting rESR repression both directly through promoter localization and indirectly by affecting Dot6 levels/activation (see Discussion). We note that Msn2/4 had a broader role in gene repression beyond Dot6 targets, consistent with previous evidence from our lab (Chasman et al., 2014). Thus, the lack of Msn2/4 has wide ranging effects on the transcriptomic response to salt (see Discussion).

A remaining question was the effect of Msn2/4 on *DOT6* mRNA abundance, given that Msn2/4 bind the *DOT6* promoter during multiple stresses (Brodsky et al., 2020; Elfving et al., 2014; Huebert et al., 2012; Kuang et al., 2017; Ni et al., 2009) and are required for normal Dot6 protein abundance (**Fig 4**). We did not see a significant expression difference in the absence of stress in bulk cultures, for *DOT6* or *TOD6* (FDR > 0.05). However, after salt stress the *msn2Δmsn4Δ* mutant showed consistently lower levels of *DOT6* mRNA compared to wild-type cells (**Fig 6**). Despite some variation in the time courses, the wild-type cells slightly and transiently repressed *DOT6* levels – cells without *MSN2/4* showed substantially stronger repression of *DOT6* mRNA. We observe a similar trend in response to H_2_O_2_ treatment (Huebert et al., 2012), heat shock (Gasch et al., 2000), and another study of NaCl stress (Chasman et al., 2014). Together with the direct binding of Msn2/4 to the *DOT6* promoter during stress (**Fig EV3**) these results indicates that Msn2/4 induction counteracts reduced expression of *DOT6* mRNA during stress, and likely also during stochastic bursts of ESR induction before the addition of stress (Bergen et al., 2022), leading to reduced protein abundance in the cell (see Discussion).

**Figure 6.**
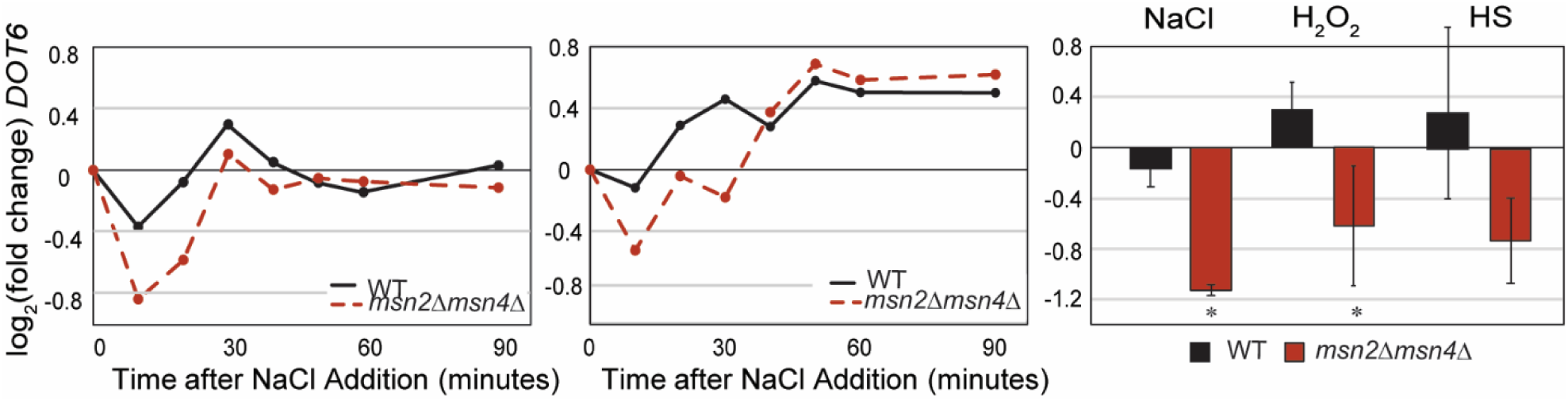
Msn2/4 are required to maintain *DOT6* mRNA levels after salt stress. **A)** Relative log_2_(fold change) in *DOT6* mRNA abundance at indicated time points compared to unstressed cells in wild-type (black) and *msn2Δmsn4Δ* cells (red) in replicate 1 (left) and 2 (right) NaCl time courses. **B)** Average and standard deviation (n=3-5) of strain responses to 30 min of 0.7M NaCl (Chasman et al., 2014), 30 min of 0.4 mM H_2_O_2_ (Huebert et al., 2012), or 20 min after a 25°C to 37°C heat shock (Gasch et al., 2000). *, p< 0.05, two-tailed T-test.

## DISCUSSION

While it was well appreciated that stressed yeast mount the common ESR response, understanding of the separable roles of iESR induction and rESR repression has remained incomplete. Our results for the first time decouple these responses to quantify the importance of resource reallocation and show that it is hard-wired to enable the induced component of the stress response (**Fig 7**). Upon acute stress, Msn2/4 transcriptionally induce their targets, most likely through direct binding of upstream STRE elements enriched in the genes’ promoters. At the same time, Msn2/4 are critical to maintain Dot6 protein levels, through direct regulation at the *DOT6* promoter. Complete loss of Dot6 and its paralog leads to failed repression of RiBi targets, thereby consuming both transcriptional and translational (Ho et al., 2018) capacity that would otherwise go toward producing defense proteins. Loss of this repression produces a drag on post-stress growth rate, delayed production of defense proteins, and slowed acquisition of stress tolerance. In addition to ensuring adequate Dot6 for gene repression, we also discovered that Msn2/4 bind promoters of many stress-repressed proteins, perhaps through an alternate mechanism since these promoters are not enriched for the Msn2/4 binding sites (**Fig 5**). As a consequence, cells lacking MSN2/4 show weaker repression of many rESR genes, likely through both direct and indirect effects. Together, these results show that Msn2/4 help dictate the resource reallocation needed for its own response. That response comes at a cost to post-stress growth rate but provides a key advantage when subsequent stresses arise.

**Figure 7.**
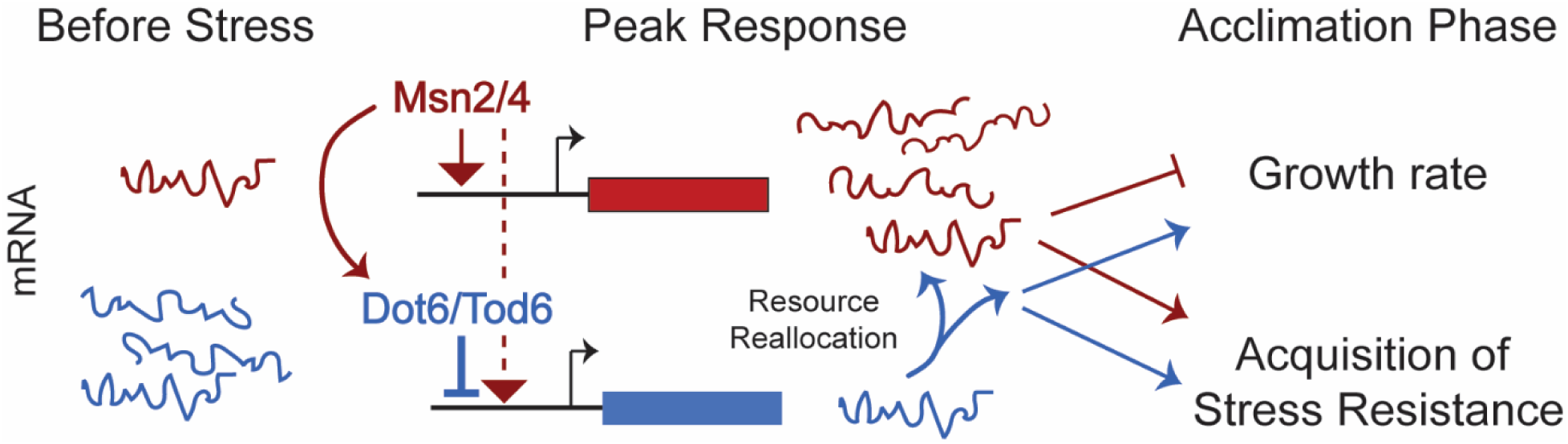
Model for Msn2-dependent resource reallocation during stress. Msn2/4 activation induces defense genes (red) and Dot6 by direct regulation, while also influencing stress-dependent repression of ribosome and growth genes (blue), both directly and indirectly by influencing Dot6 activation dynamics. Resource reallocation provided by transient rESR repression enables and accelerates the costly Msn2/4 response, promoting faster acquisition of subsequent stress tolerance, see text for details.

The insights uncovered here likely pertain to stress defense systems in other organisms as well. This includes the so-called stringent response in bacteria, where the alarmone metabolite ppGpp suppresses transcription of growth-promoting genes while supporting synthesis of proteins needed for survival (Gourse et al., 2018; M. Zhu et al., 2019). Similar themes emerge in the mammalian integrated stress response (ISR), where a host of kinases, each responding to different signs of adversity, phosphorylate eIF2α to inhibit global translation but stimulate translation of stress-responsive transcription factors that induce downstream defense targets (Costa-Mattioli & Walter, 2020; Dever et al., 2023; Harding et al., 2003; Houston et al., 2020). Results from our study in yeast can therefore reflect generalizable insights into stress responses across species.

The first is that cells alternate between objectives to balance the cost of defense versus growth. Under the conditions studied here, cells are clearly not maximizing post-stress growth rate, since they are capable of growing faster in the absence of an Msn2/4 response (**Fig 1B-C**). This adds to a growing body of evidence that maximizing growth is not a universal objective in microbes, especially during adversity (Balakrishnan et al., 2021; Basan, 2018; Basan et al., 2020; Dai et al., 2018; Ho et al., 2018; Korem Kohanim et al., 2018; Schuetz et al., 2012; M. Zhu et al., 2024). It also shows that mere abundance of ribosome-related transcripts does not predict ‘instantaneous’ growth rate, since cells with (albeit transiently) fewer RiBi transcripts grow faster after salt stress (**Fig 2** and Bergenholm et al., 2018).

Instead of maximizing growth at all costs, cells invest in preparing for impending stress at the first signs of adversity. Activation of Msn2/4 contributes to acquired stress resistance in yeast and likely other fungi (Berry et al., 2011; Berry & Gasch, 2008; Brown et al., 2014; Gasch, 2007; Liang et al., 2023). Similarly, activation of the bacterial stringent response supports acclimation to suboptimal carbon sources, by shortening the lag phase required to produce needed proteins (Balakrishnan et al., 2021; Boutte & Crosson, 2013; Gourse et al., 2018; M. Zhu & Dai, 2023). Upon a shift away from optimal carbon sources, both yeast and bacteria invest in producing enzymes for alternate sugar utilization, even when the substrate sugars of those enzymes are not present (Balakrishnan et al., 2021; Simpson-Lavy & Kupiec, 2019; Turcotte et al., 2010; Vermeersch et al., 2022). Activating these responses comes with a cost to growth rate, explaining why maximal stress tolerance is not constitutive in these organisms (Balakrishnan et al., 2021; Basan et al., 2020; M. Zhu et al., 2024; M. Zhu & Dai, 2023). But it also underscores the importance of anticipatory programs in evolution and reveals a unifying theme for fast-growing microbes like *S. cerevisiae* and *E. coli*: when times are good, cells direct resources to support maximal growth, but in response to early signs of adversity, they redirect focus to invest in the future. Similar pressures likely exist in other organisms as well.

Second, our results delineate the importance of transcriptional and translational suppression for resource reallocation during stress. The isolated role of this suppression has been hard to study in other organisms, because it is often tightly coupled with production of defense transcripts and proteins. Nonetheless, it has been suggested in *E. coli*. Cells lacking ppGpp grow fine without stress, but have a much longer lag when shifted to suboptimal conditions; conversely, ppGpp over-production, leading to stronger repression of growth-promoting genes, slows growth I the absence of stress but accelerates stress acclimation and promotes stress tolerance (M. Zhu et al., 2019, 2024; M. Zhu & Dai, 2023). Furthermore, over-production of unnecessary proteins slows cell growth rate and stress acclimation, supporting the notion that a tax on protein-synthesis capacity is suboptimal (Balakrishnan et al., 2021; Basan et al., 2020). The intimate coupling of stress-induced and -repressed responses across organisms mounting common stress responses underscores that coregulated resource reallocation is a unifying principle.

A remaining question had been how this resource reallocation is regulated in yeast. Our results show for the first time that it is in part hardwired into the ESR program (**Fig 7**). Msn2/4 maintain Dot6 protein levels through direct regulation (**Fig 4**, **6, EV3**), bind many Dot6 and other rESR promoters during stress (**Fig 5**), and ensure full rESR repression through these direct and likely also indirect mechanisms. Together through these mechanisms, these results show that Msn2/4 activity helps to orchestrate the resource-reallocation needed for its own response. But a remaining mystery is how yeast sense their internal system to set the balance between growth-promoting and stress-defending programs. *S. cerevisiae* does not utilize ppGpp, which in *E. coli* directly senses translational flux at individual ribosomes (Wu et al., 2022). In yeast, PKA and TOR may play a role, since they respond to quality nutrients to promote growth at the expense of defense (González & Hall, 2017; Kocik & Gasch, 2022). Even in an organism as well studied as *S. cerevisiae*, these mysteries await further investigation.

## METHODS

### Strains and Growth conditions

*Saccharomyces cerevisiae* strains of the BY4741 background used in this study are listed in Table 1. All strains were grown in Low Fluorescent Media (LFM) as previously described in Bergen et al., 2022 (0.17% yeast nitrogen base without ammonium sulfate, folic acid, or riboflavin; 0.5% ammonium sulfate; 0.2% complete amino acid supplement, and 2% glucose). Strain AGY2046 (*msn2Δmsn4Δ*) was generated by replacing *MSN4* in the BY4741 *msn2::KANMX* strain (Open Biosystems) with the hygromycin-MX cassette via homologous recombination and validated using diagnostic PCRs. The remaining strains were generated through genetic crosses as listed below, dissection of haploid spores, and selection of spores with appropriate markers. Gene deletions were verified by diagnostic PCR and fluorescent microscopy when appropriate. Crosses were used to generate RAKY41 (AGY2046 x AGY5), RAKY43 (AGY594 x RAKY41), RAKY35 (AGY1328 x AGY5), RAKY50 (AGY1328 x RAKY47), RAKY51 (AGY1328 x RAKY41), RAKY53 (AGY594 x RAKY35), and RAKY65 (RAKY53 x RAKY47).

**Table 1.**
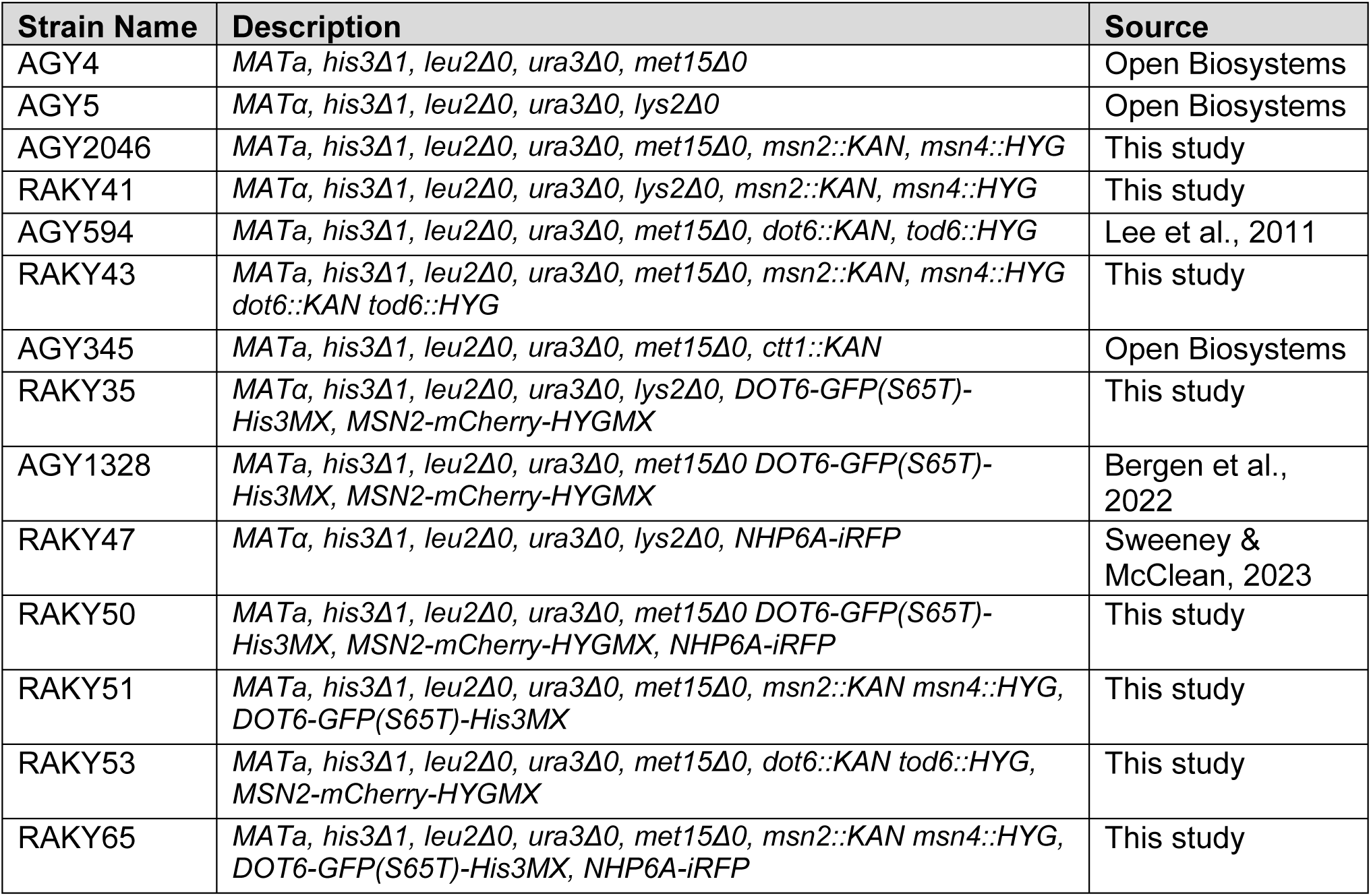
Strains used in this study.

### Liquid growth curves

Liquid cultures for growth rate assessment were inoculated in test tubes from an overnight culture grown ∼12 hours in LFM and grown for at least 4.5 hours to a starting optical density at 600 nm (OD_600_) of ∼0.1 before measurements were taken every 15 minutes. NaCl was added to 0.7 M NaCl. Growth rates were calculated by fitting exponential curves to data from time points spanning 75 minutes to 225 minutes after NaCl was added. Mutants were grown side-by-side with wild-type cultures, with paired replicates done on separate days allowing paired statistical analysis.

### Acquired Stress Resistance Experiments

See **Fig 2A** for schematic of this protocol. Cultures were grown in LFM in flasks at 30°C in a shaking incubator for at least 15 hours to a starting OD_600_ ∼0.3 – 0.4. An aliquot of unstressed cells (0 min) was removed and then NaCl was added to a final concentration of 0.7M. At various timepoints following the addition of NaCl (10, 20, 30, 40, 50, 60, 90, and 120 minutes) an aliquot of culture was retrieved, cells collected by brief centrifugation, and resuspended in fresh LFM without NaCl to an OD_600_ of 0.6. Cells were subsequently 3-fold diluted into 96-well plates containing LFM or LFM plus one of 11 doses of H_2_O_2_ (spanning from 0 to 20 mM final concentration of H_2_O_2_). Cells were incubated for 2 hours at 30°C in a shaking incubator, then a 200-fold dilution of each culture was spotted on YPD agar plates (1% yeast extract, 2% peptone, 2% glucose, and 2% agar). Plates were grown ∼48 hours at 30°C, then viability at each dose of H_2_O_2_ “secondary” stress was scored visually on a four-point scale: 100% (3), 50-100% (2), 10-50% (1), and 0% (0) survival compared to the wild-type cells treated with NaCl but no H_2_O_2_. A single H_2_O_2_ survival score was calculated for each time point as the sum of scores across the 11 different doses of H_2_O_2_. Each mutant was compared to wild-type culture grown side-by-side on each day, with 3 biological replicates for most strains except the *dot6Δtod6Δ*, which was done with 6 replicates for added statistical power.

### Microscopy and Image Analysis

Time-lapse microscopy was performed using an FCS2 chamber (Bioptechs Inc, Butler, Pennsylvania). Data collection and analyses were conducted as previously described in Bergen, Kocik et al. 2022 (Bergen et al., 2022), with the following changes. Each mutant was grown to mid-log phase in LFM media in a flask and then mixed within the microfluidic chamber at a 50:50 ratio with the iRFP-tagged wild-type strain. Media flow was switched from LFM to LFM + 0.7M NaCl after T12, as previously described. GFP and mCherry signal was recorded at each time point before and after NaCl treatment as previously described. To distinguish wild-type from mutant cells in mixed cultures, Principal component analysis of cells was performed based on the iRFP signal across timepoints T1-T40, using R Statistical Software (R version 4.3.1). This analysis led to clear dichotomy of cell types that also correlated with presence of both GFP and mCherry signal, where mutant cells showed no iRFP and loss of either GFP or mCherry signal according to the strain.

Dot6-GFP and Msn2-mCherry phenotypes were also determined as previously described, including fraction of nuclear Dot6-GFP and Msn2-mCherry signal (“nuclear/cytoplasmic ratio”, defined as the average signal of the top 5% of pixels divided by the median of all pixels), acute stress peak height, and area under the curve of fraction of nuclear signal across pre-stress or post-stress timepoints (Bergen et al., 2022). Acute stress peak height as shown in **Figs 3-4** was calculated as the maximum nuclear localization score just after NaCl addition (T13 – T20) minus the minimum fraction of nuclear signal just before salt was added (T11 – T13). Dot6 abundance was measured based on the median Dot6-GFP signal in each cell; the average signal before NaCl treatment (T1 – T12) or after (T20 – T36) is shown in **Fig 4C**. Cells shown in **Fig 4D** were defined as those with a similar level of Dot6-GFP signal (values between 635 to 650 signal intensity), such that the mutant and wild-type signal were not different (Wilcoxin rank-sum test p > 0.05).

Cell clustering in **Figs 3-4** was performed based on the population median (i.e. each column) of GFP or mCherry nuclear signal using Gene Cluster 3.0 (Eisen et al., 1998) and visualized using Java TreeView version 1.2.0 (Saldanha, 2004). The fraction of nuclear signal shown on the left was added after clustering for display.

### RNA sequencing

RNA-seq was performed using total RNA isolated from log-phase cultures in response to NaCl. Cultures were grown in LFM in flasks at 30°C in a shaking incubator for at least 15 hours to a starting OD_600_ of mid log phase. 5 mL of culture was harvested via centrifugation at 3000 RPM for 3 minutes, flash frozen in liquid nitrogen and stored at -80°C. Total RNA was extracted using hot phenol lysis (Gasch, 2002b) and purified using the RNeasy MinElute Cleanup Kit (QIAGEN, Hilden, Germany). rRNA depletion was performed with the EPiCenter Ribo-Zero Magnetic Gold Kit (Yeast) RevA kit (Illumina Inc, San Diego, CA). RNA-seq libraries were prepared using a TruSeq Stranded Total RNA kit (Illumina), and PCR purified using AMPure XP beads (Beckman Coulter, Indianapolis, IN). Paired-end sequencing was performed on an Illumina NovaSeq 6000 sequencer (Illumina).

RNA-seq reads were processed using Trimmomatric version 0.39 (Bolger et al., 2014) and mapped to the S288c genome using Bowtie 2 version 2.4.4 (Langmead & Salzberg, 2012). Read counts for each gene were calculated using HTSeq version 0.6.0 (Anders et al., 2015). Raw data can be found in the NIH GEO database Accession GSE283327. Differentially expressed genes were identified at each timepoint using a glm model in edgeR version 4.3.2 using (TMM) normalization (Robinson & Oshlack, 2010) with significance at <0.05 Benjamini and Hochberg false discovery rate (FDR) (Benjamini & Hochberg, 1995). Genes significant in at least two time points were considered for analysis. Cells with an induction (or repression) defect were defined if the gene was induced (or repressed) in a majority of time points in the wild-type cells and showed a smaller log_2_(fold change) in the corresponding mutant.

Genes whose promoters are bound by Msn2 and/or Msn4 were identified using YEASTRACT considering only evidence of direct binding (accessed on 09-23-2024) (Teixeira et al., 2023). Genes with upstream STRE consensus, CCCCT, or GATGAG were identified within 1000 bp upstream of genes using YEASTRACT (accessed on 09-25-2024) (Teixeira et al., 2023). Hypergeometric tests were performed using R Statistical Software (R version 4.3.1).

## ACKNOWLEDGEMENTS

We thank M. McClean and the McClean lab for reagents and microfluidics support, James Hose for help with RNA-sequencing, Michael Place for computational assistance, and members of the Gasch Lab for constructive feedback. This work was supported by NIH grant R01GM14975 to APG.

---

**Expanded View Figure 1.**
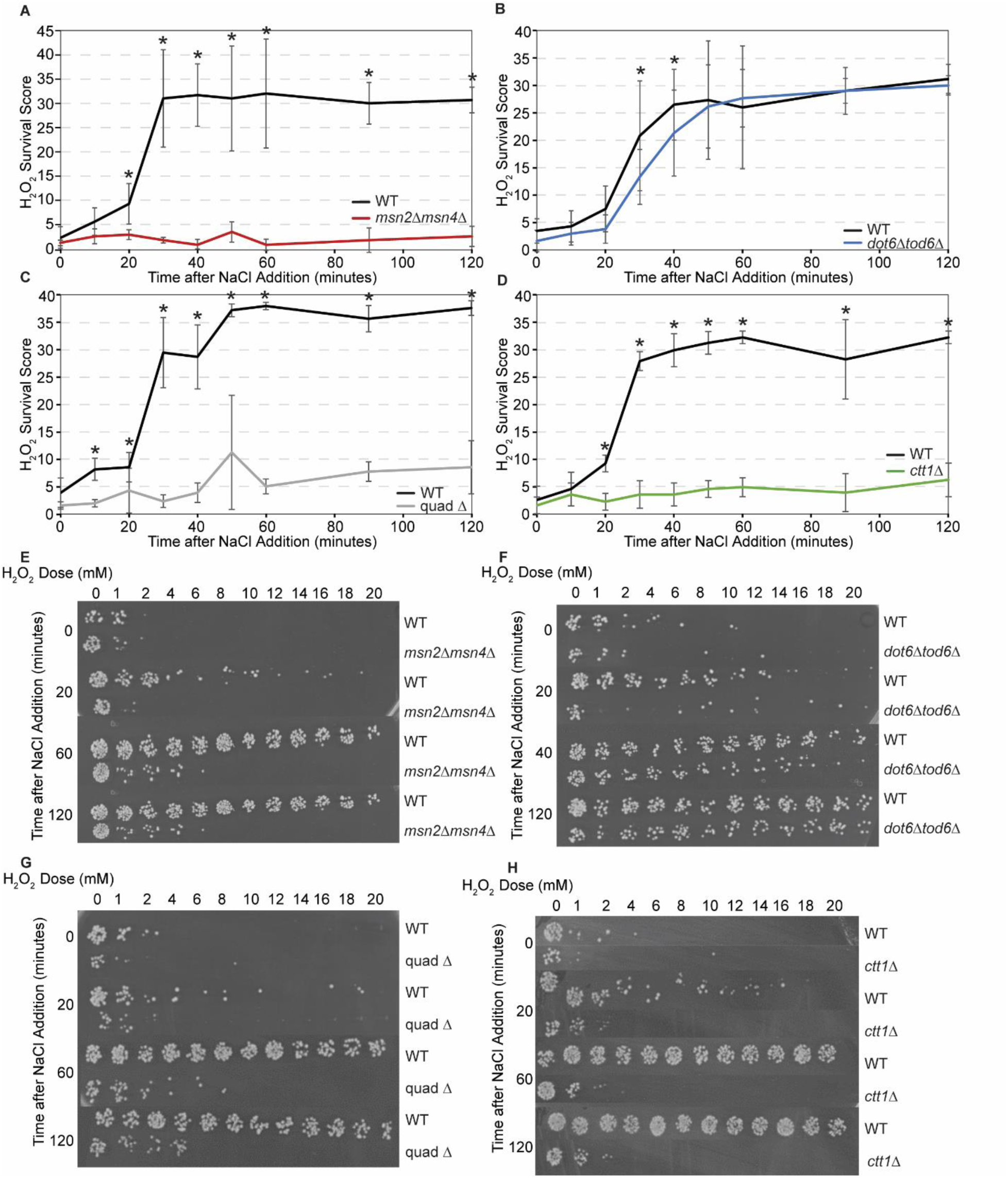
Msn2/4 and Dot6/Tod6 responses are important for acquired stress resistance. Related to **Fig 2**. **A-D)** The average change in H_2_O_2_ survival scores for wild-type (black), *msn2Δmsn4Δ* (red), *dot6Δtod6Δ* (blue), quad Δ (grey), and *ctt1*Δ (green) cells +/- 1 standard deviation, as shown in Figure 2. Colored lines are as shown in Figure 2, along with the paired wild-type culture done side-by-side each mutant. **E-H)** Representative images of cell viability across doses of H_2_O_2_ and time used to calculate H_2_O_2_ survival scores.

**Expanded View Figure 2.**
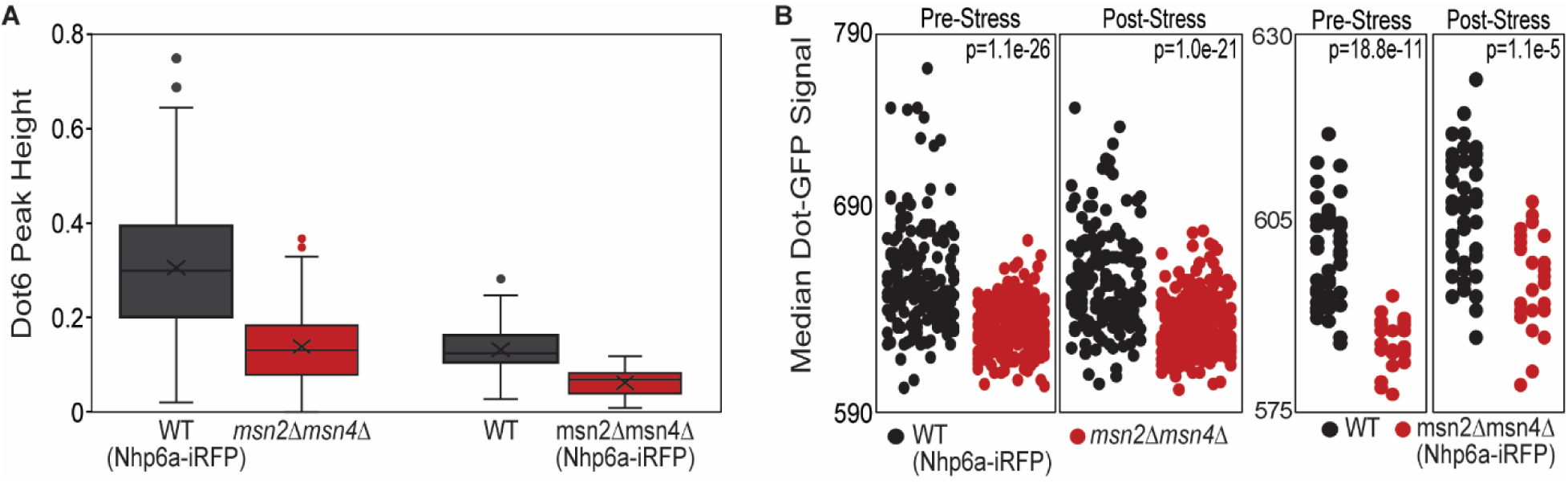
Nuclear iRFP does not affect Dot6-GFP signal. One consideration was if iRFP expressed in one strain affected GFP signal within the same strain. To ensure that our results in Fig 4 were not due to unanticipated effects of iRFP, we generated a new set of strains in which the *msn2Δmsn4Δ* cells, rather than the wild-type, carried the distinguishing iRFP signal. We found that the trends discussed in the main text were not affected by which strain carried the iRFP marker. **A)** Distribution of Dot6 acute stress peak height across wild-type and *msn2Δmsn4Δ* cells when the WT carried expressed Nhp6a-iRFP (left) or when the *msn2Δmsn4Δ* strain expressed Nhp6a-iRFP (right). Despite some difference in signal for experiments done with different laser power, *msn2Δmsn4Δ* cells showed weaker Dot6-GFP nuclear translocation signal in both sets of experiments (p=8.8e-11, left, p=1.1e-5, right, Wilcoxon rank-sum test). **B)** Distribution of median Dot6-GFP signal within the cells, scored before (0-72 min) or after (120-216 min) NaCl treatment for wild-type cells and *msn2Δmsn4Δ* cells as described in A). p, Wilcoxon rank-sum test.

**Expanded View Figure 3.**
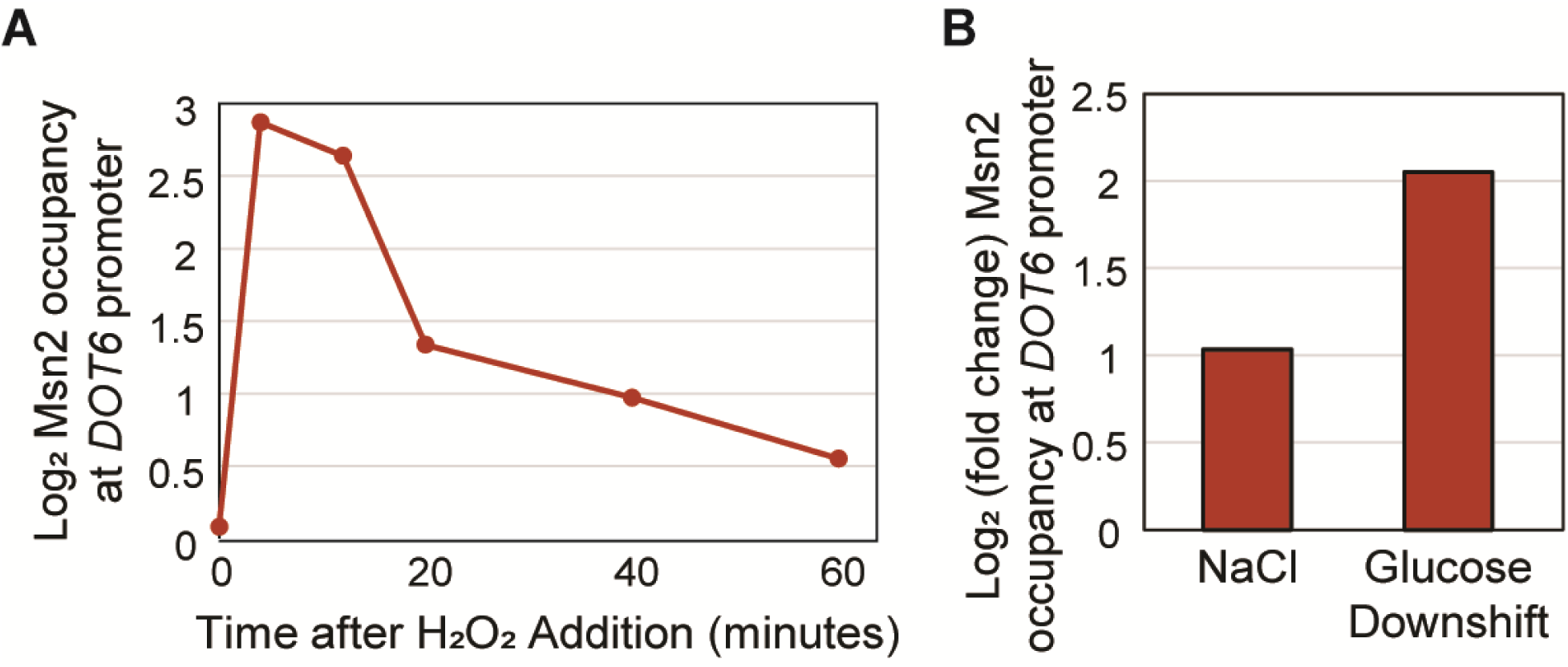
Msn2 binds *DOT6* promoter under various stresses. **A)** Log_2_ enrichment of Msn2 occupancy relative to the whole-cell extract at the *DOT6* promoter in response to 0.4 mM H_2_O_2_ (from Huebert et al., 2012). **B)** Log_2_ (fold change) of Msn2 occupancy at the *DOT6* promoter (ranging from 0 to -1000 bp from Ni et al. and +/- 250 bp surrounding the Msn2 STRE element in the *DOT6* promoter) in response to 30 min of 0.6M NaCl (left, from Ni et al., 2009) or 20 min after shift from glucose to glycerol (right, from Elfving et al., 2014) compared to the corresponding measurement in unstressed cells.

**Supplemental File 1. Microfluidics single cell measurements.** The file tabs includes microscopy measurements for A) mixed wild-type and *dot6Δtod6Δ* cells, B) mixed wild-type and *msn2Δmsn4Δ* cells, or C) mixed wild-type (without Nhp6a-iRFP) and *msn2Δmsn4Δ* cells (with Nhp6a-iRFP). Details on values are listed in D) README tab.

**Supplemental File 2. RNA-seq data.** The file tabs contain A) log_2_(fold change) measurements for all genes and all strains, and FDR values from edgeR analysis for B) *dot6Δtod6Δ* or C) *msn2Δmsn4Δ* compared to wild-type cells at each time point. Details are included in tab D) README.

